# Intergenerational shifts in innate odour preferences upon odour injections in *Bicyclus anynana* butterfly larvae

**DOI:** 10.64898/2026.03.06.710244

**Authors:** Yan Ling Chua, V Gowri, Ian Z.W. Chan, Antónia Monteiro

## Abstract

How insects transmit food odour preferences acquired during the larval stage to their offspring is unknown. *Bicyclus anynana* butterfly larvae can learn to prefer a banana-smelling odour, isoamyl acetate (IAA), via feeding on coated leaves, or simply via haemolymph transfusions from an IAA-fed animal, and transmit this preference to their naive offspring. Here we explore how larvae respond to different concentrations of IAA using olfaction choice tests, and how injections of different concentrations of IAA directly into the haemolymph impact odour learning and transmission of learned preferences. We find that naive larvae showed a slight preference towards low concentrations of IAA, and a slight avoidance towards higher concentrations. Injections of IAA at low concentrations directly into the haemolymph led to an increase in preference for IAA, whereas higher concentrations led to an increase in avoidance. Naive offspring inherited the odour preferences of their parents. Finally, injections of IAA at different concentrations into embryos did not alter choices made by hatched larvae. We establish that the same molecule (IAA) can illicit both a preference as well as an aversive reaction when directly injected into the haemolymph, but IAA is not directly implicated in intergenerational inheritance.

## Introduction

Olfaction, or the sense of smell, is pivotal in the development and expression of various animal behaviours [1]. Chief among them is foraging, involving the detection, evaluation, and movement towards attractive food odours. Which odours are deemed attractive, however, depends on the animal’s ecology: mosquitos move towards humans by recognizing a variety of body odours including carbon dioxide [2], sharks move towards prey by recognizing amines and amino acids in the water [3], and *Bombyx mori* caterpillars move towards their host plants by recognising specific leaf volatiles from mulberry plants [4].

Odours experienced by individuals will have immediate effects on behaviour, but sometimes, these effects are also manifested in the offspring, who never experienced the odours directly [5-7]. For instance, exposing lepidopteran larvae to novel food odours leads to the development of a preference for those odours in adults [8,9]. In *Bicyclus anynana* butterflies, however, such novel larval preferences are transmitted to the next generation [10]. This ability of larvae to acquire and transmit odour preferences across a single generation could be ecologically advantageous. For instance, it could encourage offspring to immediately seek out the novel food source vetted by the parental generation [10], which would promote survival in an environment with novel food plants.

The mechanism underlying food odour preference transmission in butterflies remains unclear but its effects resemble a variant of “chemical legacy”. Chemical legacy is a concept that proposes that chemical substances in the food of larvae may be retained in the larvae’s haemolymph, persist into the adult stages, and affect adult behaviours [11]. While this concept was originally developed to explain how chemicals in larval food might affect oviposition preference in adults [11], it can be extended to hypothesize that chemicals contained in larva food in the parental generation might affect food choices in the subsequent larval generation. This could happen if chemicals in the parent’s haemolymph are transferred into the zygote via sperm or eggs, altering the odour choices of larvae that develop from those germ cells.

To test whether the haemolymph contains factors responsible for the development of an odour preference and odour preference transmission to the germ line, Gowri and Monteiro [12] performed haemolymph transfusions from odour-fed and control-fed larvae, into recipient larvae. They tested larvae odour preferences immediately after the transfusion as well as preferences of their naïve offspring, more than one week later. Odour-fed haemolymph recipients no longer had an aversion toward the novel odour, and their naive offspring demonstrated a preference for the odour as compared to controls [12]. This shows that the haemolymph contains factors that induce an odour preference in the recipients and in their offspring. These factors could be the odour molecule itself that remained in the haemolymph and acted as the molecule of inheritance once it travelled and was incorporated into the cytoplasm of the germ line.

To test this hypothesis directly, here we manipulate the levels of isoamyl acetate (IAA), a banana-smelling odour, in the haemolymph of larvae. Previous studies with IAA showed that this chemical can be either an attractant or a repellent, where valency depends on the concentration used. IAA can be attractive to *Drosophila* at low concentrations (3.98×10⁻⁶ % (v/v) to 1% (v/v) IAA diluted in odourless liquid paraffin), or repulsive at higher concentrations (>1% (v/v) IAA) [13]. Congruently, naive larvae of *B. anynana* perceive 5% (v/v) IAA (diluted in ethanol) as a repulsive odour [14]. Responses toward IAA can also be sex and species-dependent, with lower concentrations of 0.0025% (v/v) (diluted in odourless liquid paraffin) being repellent to *D. suzukki* female adults, and higher concentrations of 0.005% to 0.05% (v/v) IAA being attractive to both sexes [15]. Given this diversity of responses to levels of IAA across insects, we first examined how *B. anynana* responded to different levels of IAA as an olfactory cue (Figure 1A, B). Then, after establishing that low concentrations led to a preference, and high concentrations showed a trend towards avoidance, we directly injected IAA at different concentrations into the haemolymph of recipient larvae, to test how they changed the behaviour of larvae, and of their naive offspring, towards a constant IAA odour cue (Figure 1C). Finally, we perform microinjections of different concentrations of IAA into early embryos to test whether the presence of IAA in zygotes results in an acquired preference or aversion for IAA (Figure 1D). By directly microinjecting IAA into embryos, we aimed to mimic a potential transfer of this chemical to the zygote via sperm or egg, and test its role in shaping olfactory behaviour of a neonate larva.

**Figure 1.**
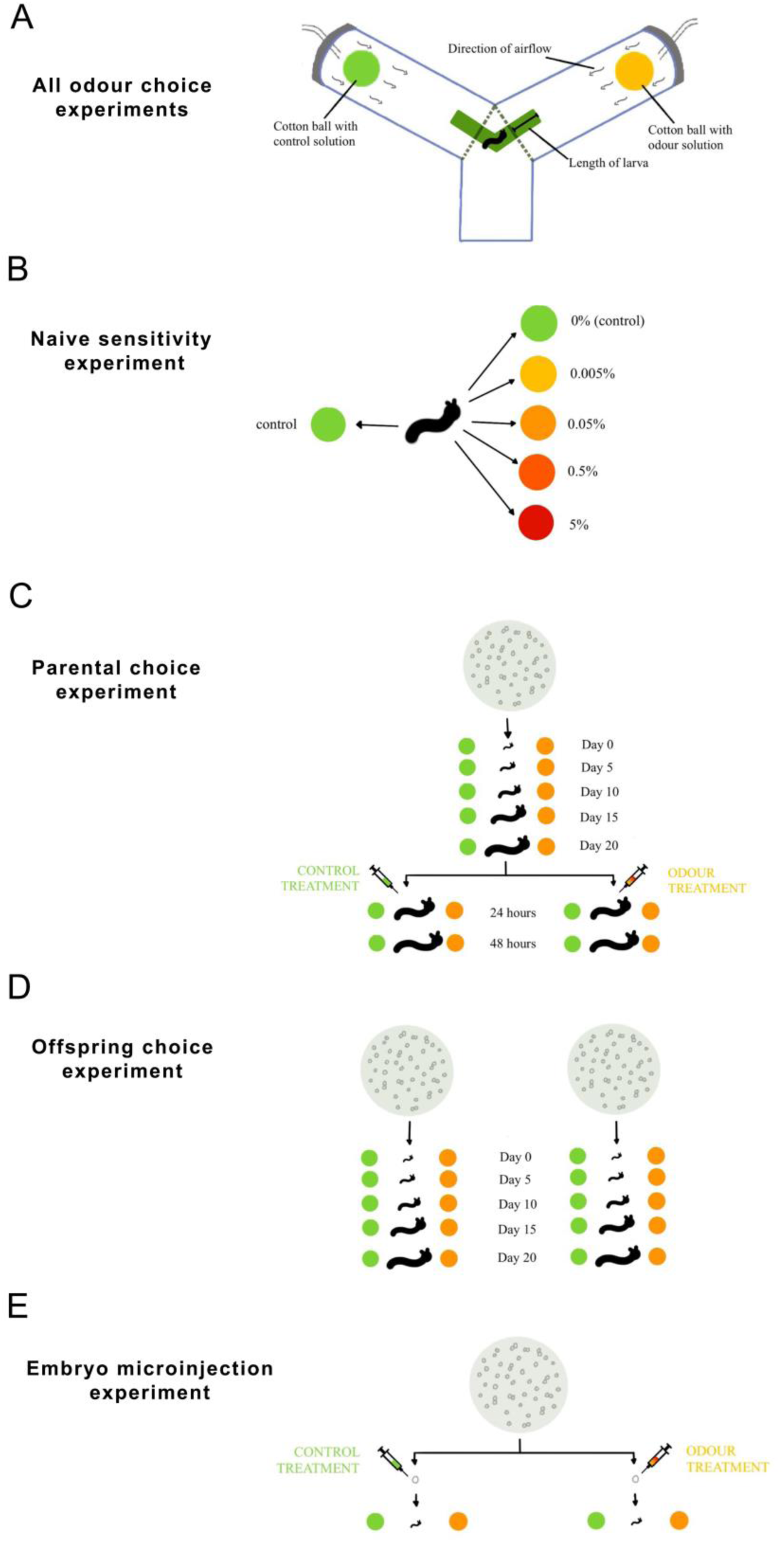
The different experiments carried out in *B. anynana* larvae.

## Materials and methods

### B. anynana husbandry

Wild-type *B. anynana* were reared in a climate-controlled room at 27 °C, 60% humidity, and 12:12-hour light:dark photoperiod. Larvae were fed with organic corn plants (*Zea mays*) that were sourced from a local farm (Greenology) in Singapore. Butterflies were fed on mashed bananas. To collect wild-type embryos, corn leaves were placed inside the adult cages for three to four hours. Eggs were removed from the leaves and collected in a Petri dish. After hatching, the larvae were reared in groups of five in plastic containers with holes for ventilation. These larvae were randomly separated and assigned to control and odour treatments for the various experiments (**Figure 1**). For all experiments, 50 larvae were used for each experimental and control group. For the egg microinjection experiment, 30 eggs were collected for each group. Larvae from each treatment were reared in the same groups of five as before and, after pupation, they were shifted to a cylindrical mesh cage for adult emergence. Larvae from the same treatment group but from subsequent batches were added to the same mesh cage after emergence to facilitate mating. All cages had fresh corn leaves for oviposition, and a Petri dish of mashed bananas on water-soaked cotton for feeding. Both the leaves and food were replaced every other day. For each experimental and control group, 50 eggs were collected in labelled Petri dishes. After hatching, choice assays were conducted on days 0, 5, 10, 15, 20.

### Olfactory Y-assay choice test

Larval odour choices were tested with a Y-tube olfactometer (see set-up in **Supplementary Figure 1)**. Different odours were placed at the tips of the arms of the Y, and carried down to the foot of the Y via air flow. Larvae were placed at the center of the Y and moved towards either arm of the Y, making a choice in the process (**Figure 1A**). Depending on the experiment, choice assays were performed on larvae on days 0, 5, 10, 15, 20, or on 5^th^ instar (pre-injection), and 24 hours and 48 hours after injection (See details below; **Figure 1**). Larvae were starved for three hours before each assay, as hungry larvae move more quickly towards odours. For each larval stage, a green coloured path, with a width of 1 cm and the length corresponding to the larvae’s body length, was drawn onto tracing paper and attached to the bottom of the flat olfactometer. The green colour acted as a stimulus for the larvae to move forward.

One ml of control and IAA solutions were dripped onto separate cotton balls. The cotton balls were placed in each arm of a Y-tube, and airflow was applied from the arms towards the foot of the Y, at a speed of 1 L min^-1^ (**Figure 1A**). Single larvae, placed in the centre of the olfactometer with a paintbrush, were given a maximum of four minutes to make a choice. Once the larva moved at least one body length towards one arm of the olfactometer, a choice for control or odour was scored. Otherwise, the larva was scored as not having made a choice. After every choice assay, ethanol was used to remove any residual body odour or silk trail that might influence the choice of subsequent individuals.

### Odour sensitivity experimental set-up

Varying concentrations of IAA solution were dripped onto the experimental cotton ball and placed on the odour side of the Y-tube, and a control solution-dripped ball was placed on the control side. Preferences of naïve 5^th^ instar larvae were scored for each IAA concentration versus the control using the choice assay (**Figure 1B**). A control group, where larvae were expected to exert a 50:50 choice to either arm of the Y-tube, had control solution-dipped balls present on each side.

### Preparation of IAA solutions

Several IAA experimental solutions were prepared from the control solution. The control solution was Ringer’s solution, and it was prepared using 7.2 g NaCl, 0.17 g CaCl_2_, 0.37 g KCl per litre of deionized water. Experimental solutions varied in IAA concentration from 0.005%, 0.05%, 0.5%, and 5% (v/v) in control solution. To make these, we used 10-fold serial dilutions of 5% of stock solution of IAA from Sigma-Aldrich (W205532, natural, 97%, Food Chemicals Codex, Food Grade) in the control solution.

### Larval and egg injections

Each 5^th^ instar larva was injected with 6 μl of IAA solution after undergoing a choice assay. The injection procedure was modified from [16] where the larvae were anaesthetised on ice for a minute and injected on a Petri dish with varying IAA concentrations using a 10 µL Hamilton 1701 removable needle (RN) syringe (Model 7653-01, Hamilton Company, USA). The needle was angled towards the head of the larva, between half to two-thirds down its length. If the larva bled when pierced, it was left to rest for at least five minutes.

Embryos (within 1hr after egg laying) were injected with different IAA concentrations using a FemtoJet 4i microinjector (Eppendorf, Singapore). We injected 30 embryos for each experimental group. Embryos were collected on a Petri dish, placed on the stereomicroscope stage at 50x magnification, and injected according to [17].

### Statistical analyses

#### Testing larvae odour choices (**Figure 1A**)

For all experiments, data from larvae that did not make a choice were removed from statistical analysis. A χ² test of goodness of fit with significance set at *P* <0.05 was used to test if the proportion of larvae choosing IAA over control solutions significantly deviated from random choice (50% - 50% choice) and was noted as a preference or aversion to IAA.

#### Testing naive odour choice of larvae (**Figure 1B**)

A generalized linear mixed-effects model with a binomial error family was fitted to model the proportion of larvae choosing IAA over control solutions in the odour sensitivity experiment. If the assumptions of the binomial GLMM were not met, the quasibinomial error family was used instead. The experimental batch that each larva came from was included as a random effect to account for variation across the ten batches used for each experimental group. This allowed for non-independence of data points that could have arisen from repeated measures, genetic relatedness, or shared experimental conditions. Larval response proportion was fitted with concentration of IAA in the cotton ball as the sole explanatory variable. Likelihood ratio tests (LRT) were used to assess the significance of concentration on larval choice.

#### Testing larvae odour choice in the parental generation throughout development across the four larval cohorts, before injection (**Figure 1C**)

A binomial GLM for larval choice was modelled with concentration (0%, 0.005%, 0.05%, 0.5%, 5% (v/v) of IAA) and days after hatching as fixed effects. Where applicable, we used LRT and a pairwise post hoc analysis using least square means with a Tukey correction to identify the pairs of experimental groups that displayed significantly different larval choices.

#### Testing the effect of injection treatment on larvae odour choice in the parental generation (**Figure 1C**)

To test for changes in the proportion of odour choice post-injection, a quasibinomial GLMM was fitted with odour concentration level injected (0%, 0.005%, 0.05%, 0.5%, 5% IAA) as a fixed effect, and time point (24 or 48 hours after injection) and batch as random effects. LRTs were used to assess the significance of all the fixed effects and their interactions. A pairwise post hoc analysis was conducted to compare odour choice between the pooled controls (larvae injected with 0% IAA solution) and experimental groups.

#### Testing the effect of IAA or control injections on offspring larvae choice (**Figure 1D**)

A quasibinomial GLMM for offspring odour choice proportions was fitted with parental odour concentration level injected (0%, 0.005%, 0.05%, 0.5%, 5% IAA) and days after hatching as fixed effects, and batch as a random effect. LRTs were used to assess the significance of all the fixed effects and their interaction. A pairwise post hoc analysis was conducted to compare odour choices between the experimental and control groups.

#### Testing the effect of various concentrations of IAA microinjected into eggs (**Figure 1E**)

A binomial GLM for larval response proportion was fitted with concentration of IAA injected as the sole explanatory variable. LRTs was used to assess the significance of concentration on larval choice.

#### Testing the effect of injecting IAA (at various concentrations) on larval mortality (**Figure 1C**)

To test if any concentration of IAA influenced larval mortality after injection in the parental generation, we used a chi-square test of independence with Yates’ continuity correction for each group versus control and significance set at *P* <0.05. The number of deaths that occurred 24 hours after injection were compared to assess any significant associations of specific concentrations with increased mortality.

All statistical analyses were performed in RStudio software (version 4.3.2) using the packages tidyr [18], dplyr [19], emmeans [20], lme4 [21], lmtest [22], DHARMa [23], and MASS [24].

## Results

### Naive larvae demonstrated an odour preference for 0.05% IAA

We first tested the odour sensitivities of 5^th^ instar larvae to different IAA concentrations. Odour concentration had a significant overall effect on larval choice χ² = 13.713, df = 4, P = 0.008). However, no individual pairwise comparison between concentrations remained significant after Tukey correction, suggesting that the behavioural response is distributed across concentrations rather than driven by a single strong contrast. A chi-squared goodness-of-fit test, however, showed that naïve 5^th^ instar larvae significantly preferred 0.05% IAA over control (N = 46, df = 1, χ² = 4.260, p = 0.039; **Figure 2; Supplementary File 1**), indicating that larvae can detect and exhibit an innate preference for IAA at this concentration. No correction for multiple testing was applied as we set to identify, a priori, a biologically effective odour stimulus. Therefore, 0.05% IAA was used in subsequent Y-tube choice assays.

**Figure 2.**
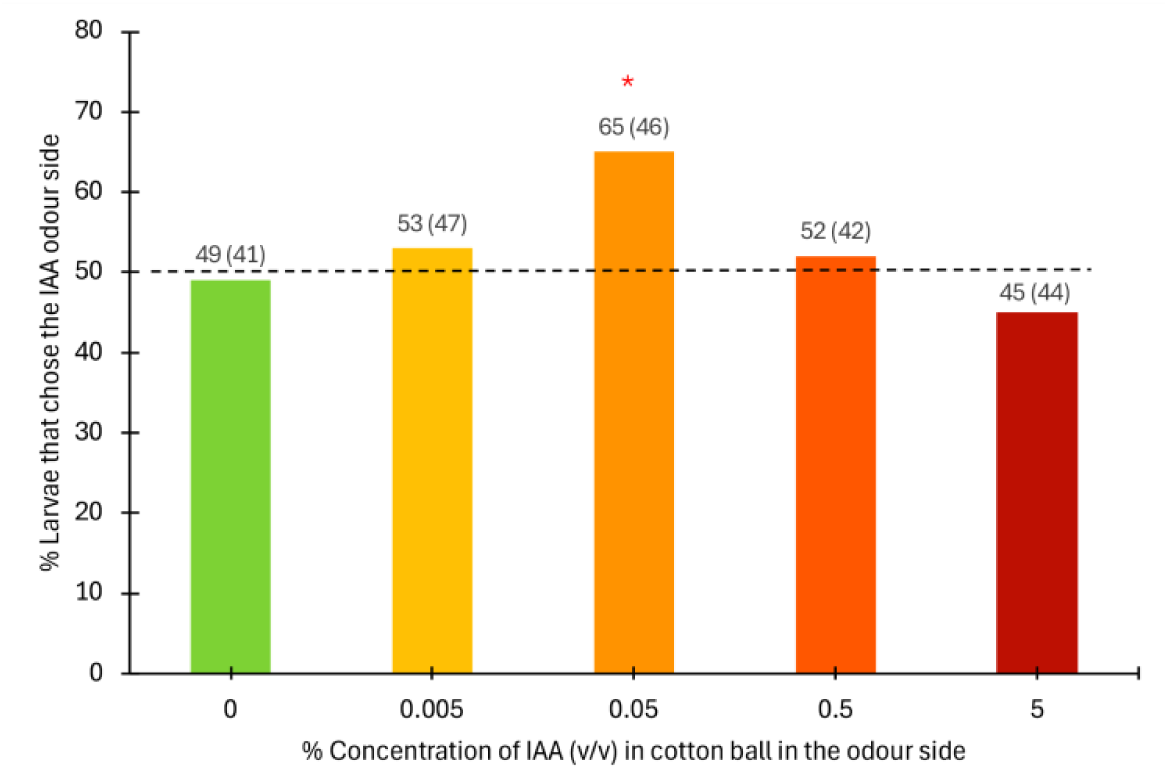
Naive 5^th^ instar larval odour choice for different concentrations of IAA. Fifty larvae were tested for each IAA concentration. Numbers above bars are read-outs from the Y-axis. The total number of larvae that made a choice is denoted in brackets. The red asterisk represents a significant preference for the odour (deviation from a random choice; Chi squared P<0.05).

### Odour injections alter odour choices in injected larvae

To test whether IAA injections (at different %) altered larval odour choices, four random cohorts of 50 larvae were first scored for their odour choices throughout development before injection.

The GLMM revealed no significant overall effect of cohort (χ² = 5.102, df = 3, P = 0.16), day (χ² = 1.596, df = 4, P = 0.81), or interaction of these two factors (χ² = 12.404, df = 12, P = 0.414) on larval odour choices before injection (**Figure 3A-D**). This indicates that larvae exhibited consistent, random odour choices regardless of their cohort or developmental stage.

**Figure 3.**
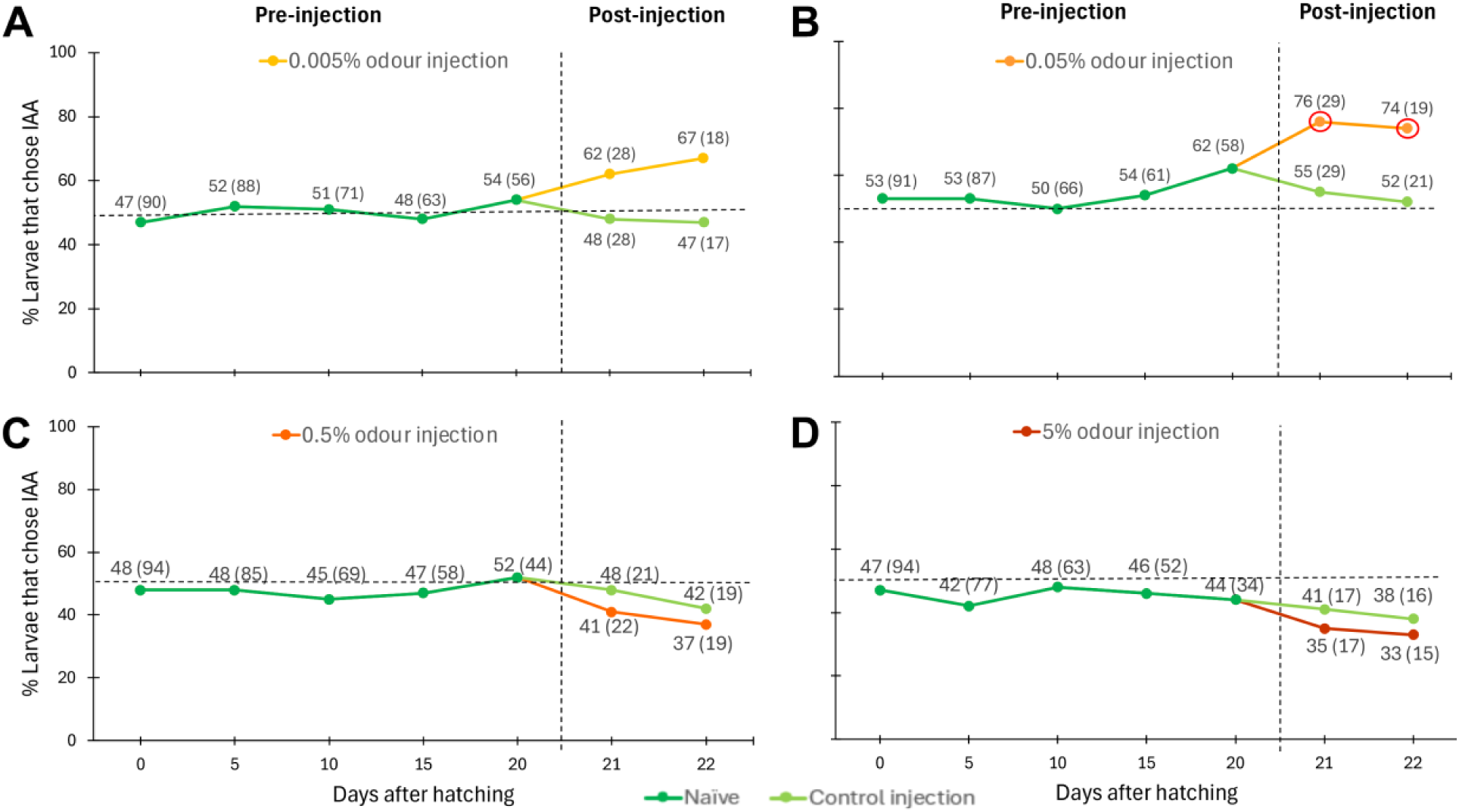
Larval odour choices during development, and after being injected with varying concentrations of IAA. At days 0, 5, 10, 15, and 20, each larva was tested for its preference for either control or 0.05% IAA odour in the Y-tube. Each cohort was then split randomly into two groups (control or experimental) and injected with control solution or (**A**) 0.005%, (**B**) 0.05%, (**C**) 0.5%, (**D**) 5% IAA concentration, respectively. The choice assay was conducted 24 hours (day 21) and 48 hours (day 22) after injection. Total cohort size for each IAA treatment group tested was 50 larvae. Numbers in graphs are read-outs from the Y-axis. The total number of larvae that made a choice is denoted in brackets. The red circles represent significant preferences (deviations from random choice; Chi squared P<0.05).

Post-injection, the GLMM revealed significant effects of odour concentration on larval preferences (χ² = 13.838, df = 4, P = 0.0078). Post hoc pairwise comparisons pointed to differences between the control vs. 0.05% IAA (P = 0.013), 0.05% vs. 0.5% IAA (P = 0.011), and 0.05% vs. 5% IAA treatments (P = 0.0067) (**Figure 3**). Larvae injected with low IAA concentrations (0.005% and 0.05%) increased preference towards IAA odour 24 hours post-injection whereas, whereas those injected with higher concentrations (0.5% and 5%), displayed slight odour avoidance (P range: 0.197 – 0.394) (**Figure 3A-D; Supplementary File 1**).

### Offspring displayed odour choices similar to their parents

To investigate whether the acquired odour preferences (or avoidances) are transmitted across generations, we mated individuals within treatment groups and scored the odour choices of their offspring immediately after hatching, and throughout development; all the while feeding them on a normal diet of corn leaves without any IAA **(Figure 4).**

**Figure 4.**
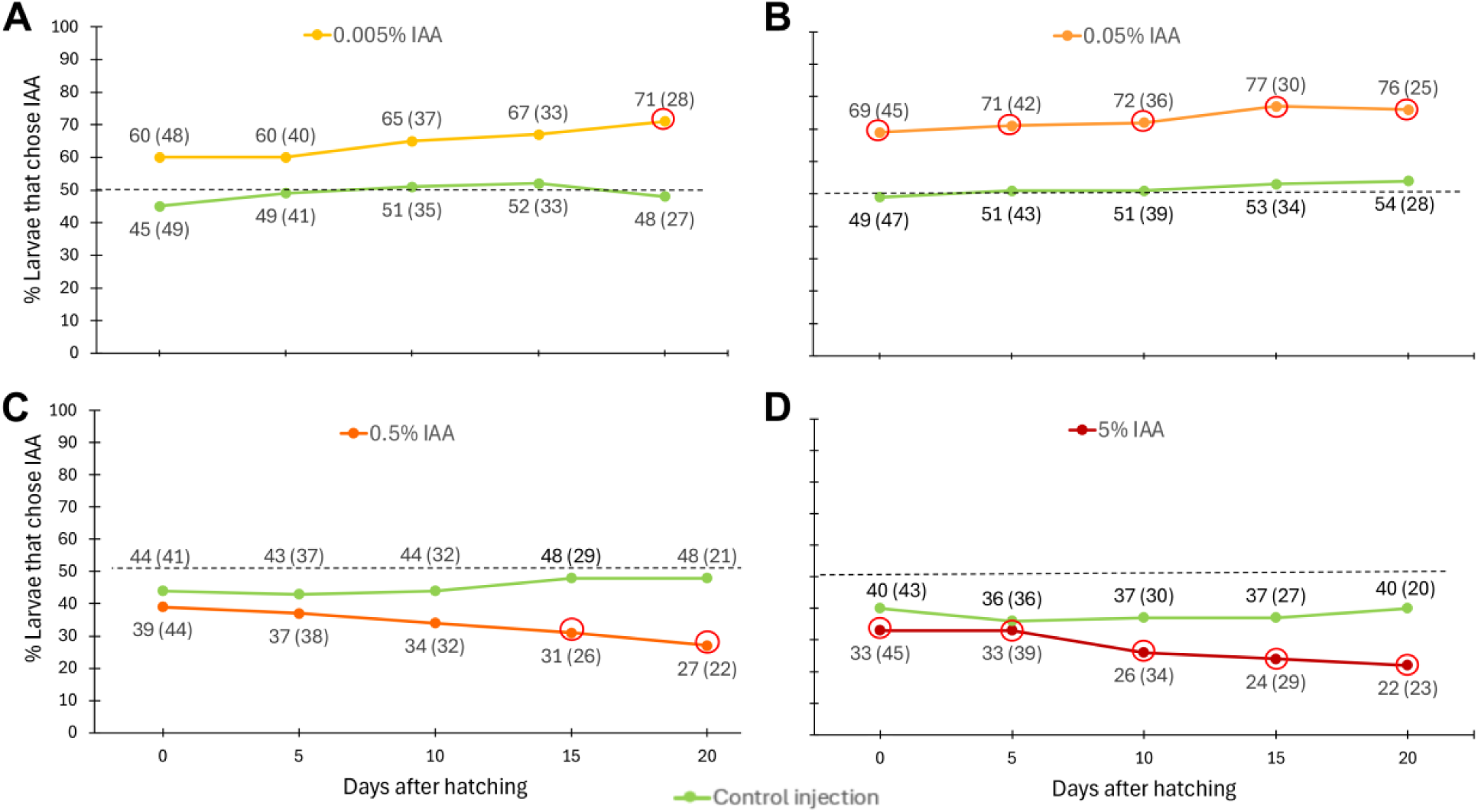
Offspring responses to either control or 0.05% IAA odour upon hatching and throughout development across the different parental odour treatments. On day 0, 5, 10, 15, and 20, each larva was tested for its choice for either control or IAA odour (0.05%). Total sample size tested per group was 50 larvae. Numbers in graphs are read-outs from the Y-axis. The total number of larvae that made a choice is denoted in brackets. The red circles represent significant preferences or aversions towards the odour (deviations from random choice; Chi squared P<0.05).

Offspring from the different parental larval treatments showed significant differences in their response to the same IAA concentration (GLMM χ² = 113.3, df = 4, P < 0.0001). Post hoc pairwise comparisons showed significant differences between the control vs. 0.005%, 0.05% and 5% IAA treatments, the 0.005% vs. 0.5% and 5% treatments, and the 0.05% vs. 0.5% and 5% treatments (all P < 0.001, except for control vs. 5% where P = 0.048; **Figure 4**). These preferences remained consistent over the twenty days of the experiment. Like the parental generation, the offspring of the larvae injected with low concentrations of IAA (0.005% and 0.05%) exhibited increased odour preference, with the effect becoming more pronounced over the course of development (**Figures 4A and 4B**; **Supplementary File 1**). The early onset and consistency of this response across developmental stages particularly in the 0.05% IAA treatment suggest that this is an optimal concentration for eliciting and maintaining an altered olfactory preference for IAA across generations. At higher concentrations, responses were reversed: the offspring of 0.5%, and particularly 5% IAA-injected parents, showed increased aversion to the odour across development (**Figures 4C and 4D**; **Supplementary File 1**), indicating that high parental IAA exposure elicits strong negative olfactory responses in offspring. The consistent directional trends observed closely mirror the observed parental behaviour. Overall, these findings highlight that the odour preference/avoidance developed by the parental generation was passed to the offspring.

### Larvae odour choice is not altered by injecting eggs with different odour concentrations

To explore whether IAA, when present in an embryo, is able to induce a preference or an avoidance behaviour in the hatchlings, we injected embryos with varying concentrations of IAA an hour after egg-laying. Larvae that hatched from these embryos displayed odour choices that deviated only slightly from those injected with control solution (0% IAA) (**Figure 5**).

**Figure 5.**
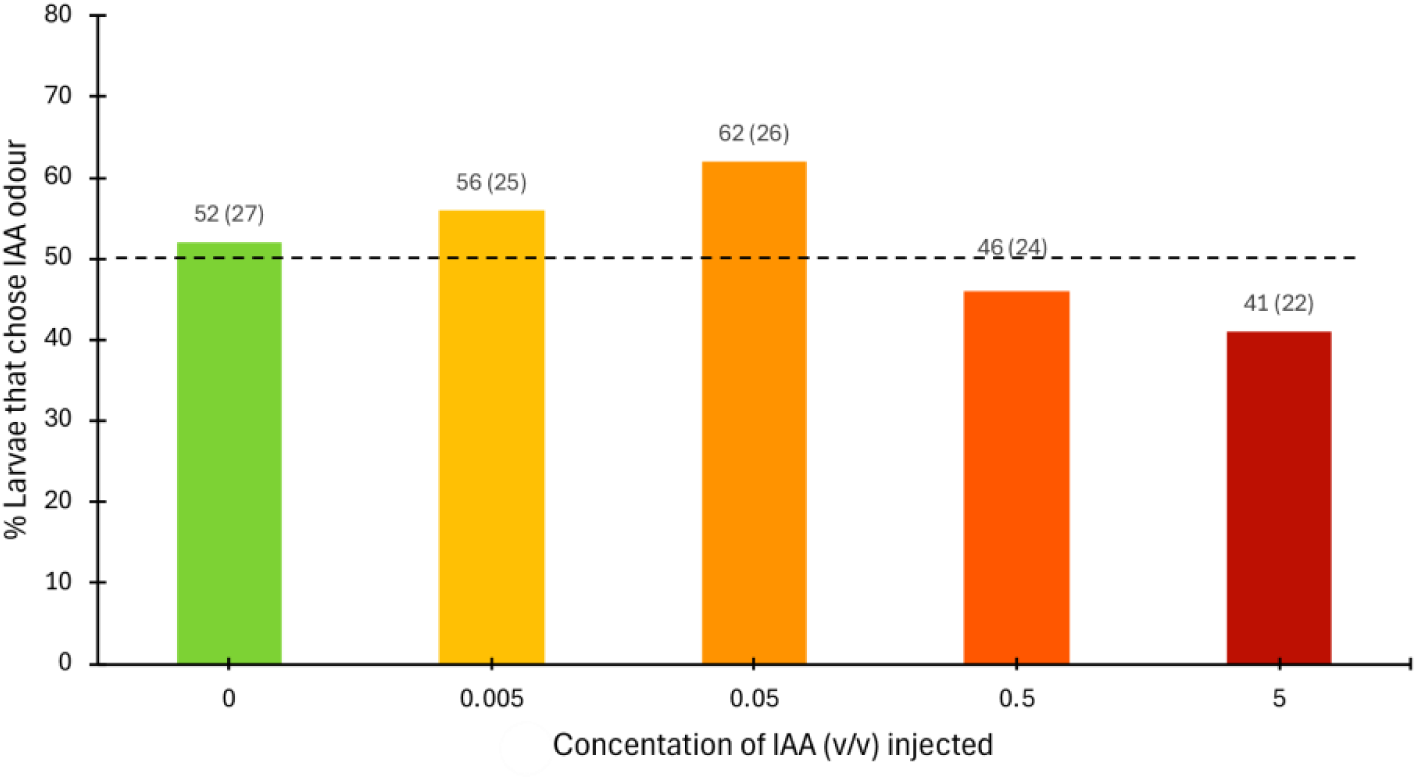
First instar larval odour choices following injections of IAA at 0%, 0.005%, 0.05%, 0.5%, 5% into 1 hr old embryos. Each larva was tested for its preference for either control solution or IAA odour (0.05% v/v) after embryonic injections with varying concentrations of IAA. Total sample size for each IAA concentration tested was 30 larvae. The total number of larvae that made a choice is denoted in brackets.

Although the GLMM revealed that there was no significant effect of concentration injected into embryos on larval odour choices (χ² = 2.5642, df = 4, P = 0.6332) (**Figure 5**), the overall trend observed parallels the patterns observed in the parental and offspring generations. Larval odour choice may reflect the concentration of odour molecules present in embryos, shaping larval behaviour and choices. However, more data is necessary to support this hypothesis further.

### Parental larval morality following IAA injection

To test whether parental larval mortality was associated with IAA concentration injected into larvae, we conducted a chi-square test of independence. Mortality increased with higher IAA concentrations in a dose-dependent manner (**Figure 6**). Chi-square tests indicated significantly higher mortality for 0.5% IAA (χ² = 22.87, df = 1, p < 0.001) and 5% IAA (χ² = 45.53, df = 1, p < 0.001) relative to control (0% IAA), with observed mortality rates of 44.2% and 75.0% compared with 6.3% in control. While the 0.005% IAA group also showed a statistically significant difference (χ² = 19.65, df = 1, p < 0.001), this very low concentration is not expected to be biologically toxic, suggesting the difference is likely due to random variation. There was no significant difference in mortality for 0.05% IAA relative to controls (χ² = 0.63, df = 1, p = 0.429). These results suggest that higher concentrations of IAA (0.5% and 5%) may have toxic effect on larvae.

**Figure 6.**
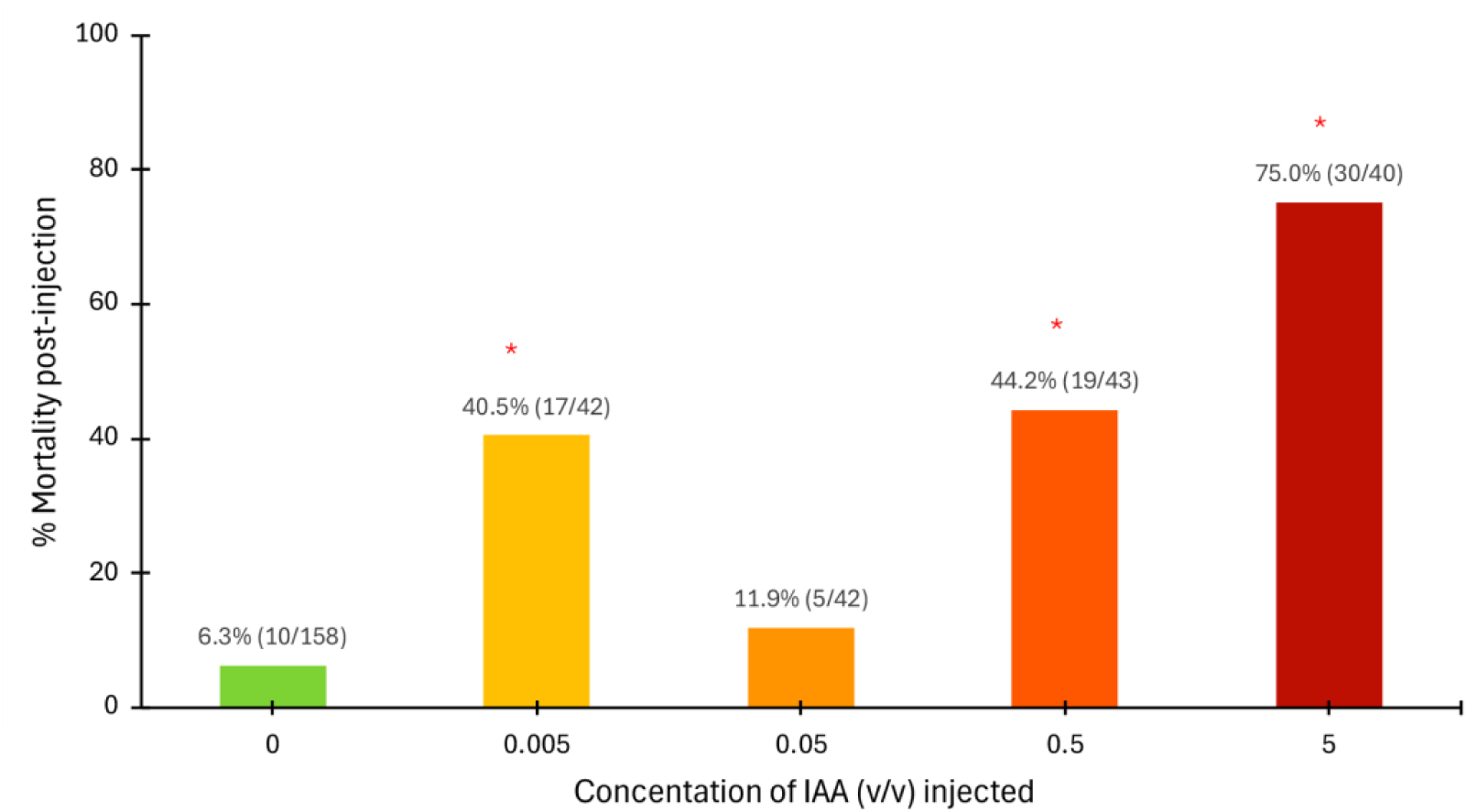
Larval survival post-injection for varying concentrations of IAA. Orange shaded bars represent larvae injected with varying concentrations of IAA (0.005%, 0.05%, 0.5%, 5% (v/v) IAA), whereas the green bar represents cohorts injected with control solution (0% IAA). The red asterisk represents a significant increase in mortality (P < 0.05) relative to controls, using chi-square statistics. Numbers in brackets represent larvae that died and total larvae injected.

## Discussion

In the current study, we showed that IAA odour molecules directly injected into the haemolymph of 4^th^ instar larvae of *B. anynana* altered odour preferences in these same larvae, one and two days later, and in their offspring, many days later, in a concentration-dependent manner. Previous experiments with haemolymph transfusions from IAA-fed and control-fed larvae into recipient larvae showed that factors in the haemolymph could make recipient naive larvae develop a preference towards IAA odours, and that these preferences were transmitted to their offspring [12]. Here we show that IAA directly injected into the haemolymph of recipients has the same effect as haemolymph transfusions, but only when concentrations are low. When concentrations are high, IAA injections lead to an avoidance behaviour in Y-tube olfactometer tests. Furthermore, we show that preferences or avoidances induced in the parents are inherited by their offspring.

The underlying mechanism for how injections of an odour into the haemolymph of an insect affect odour preferences in that same generation remains underexplored. Prior research in pigeons, turtles, and rats showed that injections of different odours into the blood of these animals was detected by odour receptors in the nose [23]. As such, it is possible that blood or haemolymph-born odours are perceived by traditional odour or gustatory receptors in sensilla on the proboscis, labial palps, and antennae, entering these cells via the blood, instead of via the air [24]. Alternatively, injected odourants may influence behaviour by acting on central processing pathways in the brain, bypassing the peripheral olfactory or gustatory systems.

Our results reveal a concentration-dependent shift in behavioural responses to IAA in *B. anynana* larvae and in their offspring. At low concentrations (0.005% and 0.05% (v/v)), a greater proportion of larvae walked towards IAA odour compared to controls, whereas at higher concentrations (0.5% and 5% (v/v)), fewer larvae walked toward IAA (**Figures 4 and 5**). This non-linear behavioural pattern parallels that of *D. melanogaster* larvae, that prefer low concentrations of IAA (diluted in odourless liquid paraffin from 3.98×10⁻⁶ % (v/v) to 1% (v/v)) but avoid it sharply when presented at higher levels [13].

Such non-linear, concentration-dependent behavioural responses to odorant concentration have been documented across diverse insect species, although the exact thresholds differ by species, life stage, and chemical identity [13]. For instance, *D. suzukii* and *D. melanogaster* respond to different minimum concentrations of IAA in both larval and adult stages [13,15]. In *D. melanogaster*, the IAA receptor Or22a exhibits a half-maximal response at a dilution of 10⁻⁴·² in paraffin oil [27]. Honeybees show a similar pattern with caffeine, which enhances memory at low concentrations but becomes aversive at >1 mM [28]. Collectively, these studies illustrate that plant-derived compounds often act as attractive cues at low doses but become deterrents or toxic at higher levels.

Our findings further suggest that *B. anynana* may possess both a lower detection threshold and an upper toxicity limit for IAA. Although specific IAA receptors have not been identified in this species, the behavioural switch we observed likely reflects sensory thresholds like those described in other insects. Notably, the 5% IAA (v/v) treatment produced a significant increase in mortality, indicating that this concentration may exceed a physiological tolerance limit. Comparable dose-dependent toxicity has been documented in other plant-insect systems. For instance, nicotine in floral nectar attracts bumblebees at low levels but becomes strongly deterrent and harmful at higher concentrations [29]. Thus, IAA becomes both behaviourally aversive and physiologically damaging to *B. anynana* at high concentrations.

How odour valence shifts from attraction to repellence remains poorly understood, but changes in receptor activation and neural processing depending on IAA concentration may play a key role. In *D. melanogaster* larvae, low concentrations of (E)-2-hexenal activate the Or7a receptor to drive repulsion, whereas higher concentrations activate additional receptors that override or saturate Or7a, yielding an attractive response [30]. Similarly, in *Aedes aegypti*, exposure to low concentrations of the attractive compound 1-octen-3-ol decreases transcription of its receptor Or8, suggesting that odour abundance can induce receptor downregulation and reduce odour sensitivity [31]. These examples highlight the potential for concentration-dependent receptor recruitment or modulation to explain the behavioural shifts we observed in *B. anynana*, though the underlying mechanisms require direct study.

The mechanism by which parents transmit learned odour preferences to their offspring is also unclear. One possibility is direct chemical transmission via the germline, such as through the spermatophore or egg cytoplasm [32]. Such molecules could influence the sensory or neural embryonic development of naïve offspring. However, many plant-derived esters, including IAA, are subject to enzymatic biotransformation in insects [33-35], and while IAA degradation via esterases has been shown in vitro, convincing in vivo evidence is lacking. Whether *B. anynana* metabolises or retains IAA at levels relevant for transgenerational signalling therefore remains an open question.

Our injections of IAA directly into embryos did not provide strong evidence that the molecule alone transmits odour preferences. Although slightly more larvae chose IAA when embryos were injected with low IAA concentrations (0.005% and 0.05% (v/v)), and fewer chose it when injected with high concentrations (0.5% and 5% (v/v)), these trends were small and statistically non-significant (**Figure 5**). Larger sample sizes and tests of wider concentration ranges are needed to determine whether IAA can drive learned behavioural transmission. Chemical analyses, such as quantifying IAA levels in larval haemolymph and eggs via gas chromatography-mass spectrometry (GC-MS) analysis would also clarify whether injected IAA disperses systemically or persists long enough for transgenerational transfer. Such studies would help elucidate both the chemical and neural mechanisms underlying odour learning and inheritance in *B. anynana*.

## Conclusions

Our findings contribute to a growing body of evidence suggesting that insect olfactory preferences are plastic and can be transmitted to the next generation [7,10,12,14,36]. Early experience with novel odours can override innate aversions [10], and this could be advantageous for larvae if they find themselves on novel host plants that have novel chemical signatures (via an oviposition mistake by the mother, for instance). This plastic behaviour can potentially facilitate the colonization of novel host plants and relieve individuals from intraspecific competition. Our study also raises intriguing questions about how internal haemolymph-born exposure to chemicals alters olfactory behaviour. Understanding these mechanisms in future may offer valuable insight into the evolution of sensory systems and behavioural adaptation in insects.

## Acknowledgements

This research was supported by National Research Foundation (NRF) Singapore, under its Investigatorship programme (NRF-NRFI05-2019-0006 Award), a Yale-NUS PhD scholarship to V.G., and the Ministry of Education (MOE) Singapore award MOE2018-T2-1-092.

## Supplementary Figures

**Supplementary Figure 1.**
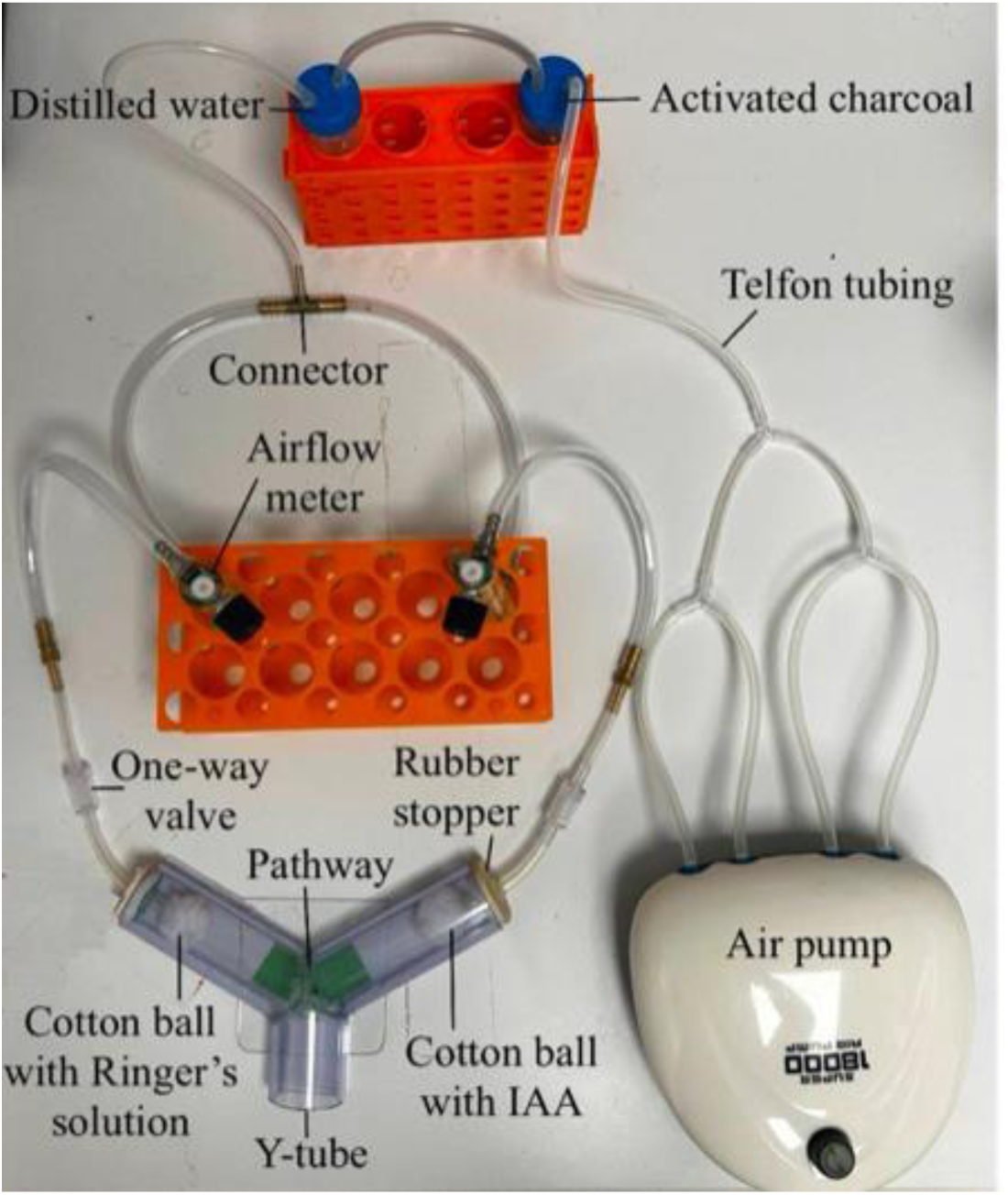
Y-assay olfactory experimental set-up to test larval odour choices. Air was pumped through activated charcoal and distilled water for purification. The airflow of 1 L min^-1^ was adjusted using two airflow meters connected via Teflon tubing to the Y-tube. Cotton ball dripped with control solution or IAA was placed at the two ends of the Y-tube as the odour source. The length of the foot of the pathway was adjusted according to the average length of the larvae that were tested. The colouration provided a stimulus to encourage the larva to move towards either of the Y-tube arms containing the odour source.

